# Computational Analysis of Lifespan Experiment Reproducibility

**DOI:** 10.1101/107417

**Authors:** Michael Petrascheck, Dana L. Miller

## Abstract

Independent reproducibility is essential to the generation of scientific knowledge. Optimizing experimental protocols to ensure reproducibility is an important aspect of scientific work. Genetic or pharmacological lifespan extensions are generally small compared to the inherent variability in mean lifespan even in isogenic populations housed under identical conditions. This variability makes reproducible detection of small but real effects experimentally challenging. In this study, we aimed to determine the reproducibility of *C. elegans* lifespan measurements under ideal conditions, in the absence of methodological errors or environmental or genetic background influences. To accomplish this, we generated a parametric model of *C. elegans* lifespan based on data collected from 5,026 wild-type N2 animals. We use this model to predict how different experimental practices, effect sizes, number of animals, and how different ‘shapes’ of survival curves affect the ability to reproduce real longevity effects. We find that the chances of reproducing real but small effects are exceedingly low and would require substantially more animals than are commonly used. Our results indicate that many lifespan studies are underpowered to detect reported changes and that, as a consequence, stochastic variation alone can account for many failures to reproduce longevity results. As a remedy, we provide power of detection tables that can be used as guidelines to plan experiments with statistical power to reliably detect real changes in lifespan and limit spurious false positive results. These considerations will improve best-practices in designing lifespan experiment to increase reproducibility.

## Introduction

Over the last few years, science has been plagued by a reproducibility crisis (Editors 2005; Ioannidis 2005; Baker 2016). This crisis has also taken root in the aging research community, with several high-profile controversies regarding lifespan extensions. Frequently cited reasons for the failure of a result to reproduce are substandard technical ability, lack of attention to detail, failure to control environmental factors or that the initial positive result was a statistical outlier that was never real in the first place. One way to address these reproducibility problems would be to list the numerous controversies and to attempt to identify the individual underlying causes and to provide a possible explanation (see note #1). This would be a long and arduous task resulting in largely speculative explanation and provide little in terms to resolve future controversies. An alternative way would be to assume that these controversies arise mostly through honest disputes of scientists standing by their results. If so, their frequency would suggest an underlying technical problem with standard practices in the field that foster such disputes. We decided to take the alternative way and to ask how reproducible lifespan experiments are under ideal conditions, *in silico*, allowing to control every environmental and technical aspect.

Statistical analysis provides powerful tools to increase reproducibility. Statistics provides a metric on how likely it is to observe a result by chance (Krzywinski & Altman 2013). In statistical analysis of experiments that measure the effect of pharmacological, genetic, or environmental factors on lifespan, the null hypothesis, H_0_, states that the perturbation has no effect on the lifespan. This hypothesis is commonly tested in lifespan experiments by comparing Kaplan-Meier survival estimates and log-rank tests (Ziehm & Thornton 2013). The two advantages of the Kaplan-Meier / logrank approach are that i) they are non-parametric and do not require the data to be distributed in a specific way (e.g. normal distribution) and ii) that they include a method to deal with censored data, assuming that the censored animals would otherwise have the same survival function of non-censored animals (Cleves *et al.* 2004). As with any type of statistical test, Kaplan-Meier analyses are subject to two types of statistical errors: Type I errors, or false positives, reject a true null hypothesis (H_0_: no extension in lifespan), whereas type II errors, or false negatives, retain the hypothesis, H_0_ when it truth it should be rejected in favor of an alternative hypothesis (H_1:_ increase in lifespan). Both errors affect reproducibility. If a result does not appear reproducible the original finding could be the result of a false positive or the failure to replicate the result could be due to a false negative result.

One important experimental consideration to minimize both false positive and false negative results is the power of detection (POD), or statistical power of a given experimental design. POD is defined as the probability to appropriately reject H_0_ in favor of the alternate hypothesis. For lifespan experiments, where H_0_ is that there is no effect on lifespan, the POD is the probability to correctly detect a true lifespan extension. Power calculations are a statistical tool to determine whether the experimental design is sufficient to detect the expected effects size. Power calculations are widely used in long term expensive mouse experiments or in clinical trial to ensure that the planned experiments have the necessary power to detect the expected effect. However, power calculations are rarely employed in experiments to measure the effects of genetic or environmental perturbations that could affect lifespan in invertebrate model organisms such as *C. elegans*.

In this study, we asked how POD is influenced by different experimental practices and how likely it is that underpowered experiments lead to scientific disputes between two groups conducting identical experiments. To address these questions, we generated a parametric model based on the Gompertz equation using lifespan data of 5,026 *C. elegans* (Johnson 1990; Ye *et al.* 2014; Kirkwood 2015). We then used this model to simulate lifespan experiments with different conditions to determine how experimental parameters affect the ability to detect lifespan increases of certain sizes. We considered two important experimental features that contribute to the workload of lifespan experiments: frequency of scoring and number of animals in each cohort. Our data show that the POD is greatly affected by the number of animals in each group, but less so by scoring frequency. We further show how inappropriately powered experiments negatively affect reproducibility. Our results make clear that current standard practices are unlikely to consistently reproducible results for real longevity effects below 20%, even under ideal conditions. We provide a series of power calculation tables to be used as general guidelines to plan and execute *C. elegans* lifespan experiments with adequate power of detection, though our approach is applicable to any organism for which a robust set of mortality data is available to derive the parameters of the Gompertz equation. Taking our findings into consideration will increase the reproducibility of lifespan measurements, thereby avoiding unproductive controversies, and help ensure the field focuses on genetic and environmental factors that have real effects on longevity in animals.

## Results and Discussion

The goal of this study was to evaluate how different experimental parameters could affect reproducibility of lifespan measurements. To accomplish this, we derived a parametric model based on the Gompertz equation that allowed us to simulate thousands of lifespan experiments *in silico*. Our model, described below, is based on experimental observation of 5,026 wild-type (N2) *C. elegans*. We then used this model to simulate lifespan data collected under different experimental parameters to evaluate how different factors influence lifespan curves and their reproducibility. The great advantage of this in silico approach is that it allowed us to precisely control sources of variability and experimental behavior; therefore, our results demonstrate the effect of each manipulation under ideal conditions.

### A parametric model to simulate lifespan data

To simulate lifespan data that share the distribution and characteristics of real experimental data, we generated a parametric model that describes *C. elegans* mortality data from the observed lifespans of 5,026 N2 control animals of one of our previous lifespan screens (Ye *et al.* 2014). The lifespan of these animals was measured in liquid media at 20ºC and resulted in an average lifespan of 21.1 days, a median of 22 days, and a maximum lifespan, defined as the longest lived 10^th^ percentile of 32 days, which are all in the range of the published literature (Kenyon *et al.* 1993). We fit these data with the Gompertz model and used regression analysis to determine the parameters G and γ (Fig. 1a, b).

**Fig. 1.**
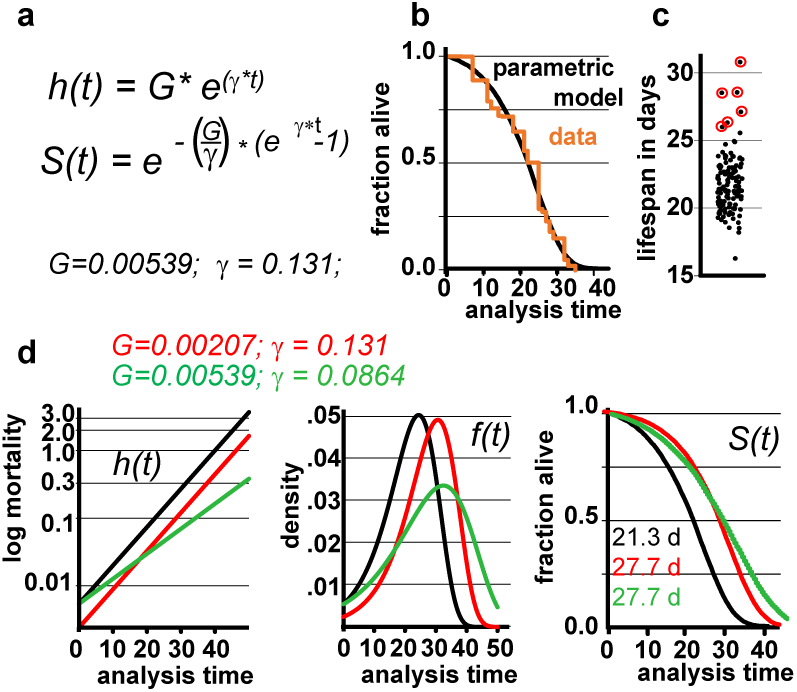
Gompertz-based Parametric Model for *C. elegans* lifespan. a) Gompertz equation, h(t), describing the mortality (hazard) function, and its corresponding survivor function, S(t). The fit parameters G and γ from wild-type lifespan data are shown in black. b) Maximum likelihood estimation was used to fit a parametric model (black) to the survival data of 5,026 N2 control animals (experimental data in orange), with a 95% CI for γ=0.127 to 0.134. c) Scatter plot showing the experimentally-measured lifespan of 127 population of ~40 animals each. The 6 populations treated with mianserin are indicated with red circles (33% increase). d) Hazard function h(t), survival time S(t) and density function f(t) plotted for wt (black) and cohorts with a +30% increase in lifespan, achieved by either modulating G (red) or γ (green).

We experimentally validated our parametric model to ensure that the underlying assumptions were sound. For this test, we took advantage of the observation that Mianserin, an atypical antidepressant, increases *C. elegans* lifespan by ~35% (Petrascheck *et al.* 2007; Ye *et al.* 2014). Mianserin extends lifespan by causing a perfect parallel shift of the lifespan curve, suggesting a modulation of the Gompertz G parameter (Kirkwood 2015; Rangaraju *et al.* 2015). We used the S(t) equation (Fig. 1a) to calculate G for Mianserin-treated animals, and then simulated 10,000 lifespan datasets consisting of a control and Mianserin-treated cohorts of 40 animals each. For each of the 10,000 datasets we ran a log-rank test to determine how likely the experimental design was to detect the lifespan extension by Mianserin. Our simulation predicted that 99% of Mianserin-treated populations should be detected by this experiment. To test this prediction, we recorded the lifespan of an additional 127 N2 populations, 4,994 animals total, of which six populations were treated with Mianserin (50µM). The experiment was conducted blind with the experimenter knowing neither the number of Mianserin-treated populations nor their identity. As predicted by our simulation, all Mianserin-treated populations were distinguished from untreated controls with P values ranging from 0.0057 to 1.6E-8 (red circles, Fig. 1c). We conclude from this test that our parametric model agrees well with real experimental data from lifespan experiments.

Knowing the values of G and γ allowed us to generate a parametric model of wild-type (N2) lifespan data by defining the density function f(t), and the survivor function S(t) based on the Gompertz equation h(t) (Fig. 1d). We have provided these equations and a description of how they relate to each other in Supplementary materials. To generate random lifespan data sets with the distributions we observed in lifespan data of the 5,026 N2 control dataset we derived the quantile function Q(u) with u being a random number between 0 and 1. The quantile function Q(u) enables Monte-Carlo simulations of lifespan experiments by creating pairs of datasets with different lifespans and to test for statistical significant differences between them busing the log-rank test (see material and methods). This Q(u) equation can be used to simulate lifespan data for any species for which the parameters G and γ have been derived from an existing dataset. To test our parametric model, we generated 10 random dataset of 5,026 animals each and compared it to the original experimental data. The simulated cohorts had a lifespan of 21.15 +/- 0.1 days (vs 21.1 days), a median lifespan of 22.0 +/- 0.1 days (vs 22 days), and a maximum lifespan defined as the longest lived 10^th^ percentile of 30.83 days +/- 0.1 vs (32 days) (Fig. 1 a, b). Thus, our model generates lifespan data in close agreement with empirical observation.

We based our simulations on the Gompertz equation because of its extensive use in the past and because it models death times reasonably well across many species (Johnson 1990; Bronikowski *et al.* 2002; Mair *et al.* 2003; Jones *et al.* 2014; Kirkwood 2015; Koopman *et al.* 2016). There are, however, some limitations to the Gompertz model. These limitations only minimally affect the results presented, but are nevertheless worth mentioning. The Gompertz model was conceived by fitting an equation to a large human mortality data set to calculate life insurance premiums (Kirkwood 2015). Insofar as the Gompertz equation is not based on a mathematical formulation of how aging works, its parameters G and γ do not necessarily describe any biological or molecular entity. However, it is common to interpret the G as a measure of initial mortality, or the mortality of individuals before the beginning of physiological decline, and γ as the “rate of aging” (Johnson 1990; Rangaraju *et al.* 2015). Although the Gompertz curve fits mortality reasonably well across lifespan, in many species the ~last 5% of the animals do not die off exponentially and therefore deviate from the Gompertz curve (Vaupel *et al.* 2004; Kirkwood 2015; Stroustrup *et al.* 2016). We cannot correct for this inaccuracy without either compromising the accuracy of mean and median lifespan or by changing the equations. These are minor issues with regards to the power of detection as the fraction of animals affected is small. However, this feature does effect the determination of maximum lifespan, as can already be observed by the somewhat larger error in our simulated data with regards to maximum lifespan compared to the actual data (30.83 vs 32 days).

For the remainder of this study, we used our experimentally-validated model to generate power of detection (POD) plots and to determine how the POD, and thus reproducibility, is affected by different experimental practices or circumstances. In these studies, each data point in the POD plots shown was derived by conducting 10,000 simulations to determine the probability of the log-rank test to detect a statistical significant difference. Whether a result is deemed significant or not is determined by setting a significance level, a, which describes how likely it is for each result to occur by chance. We present data with α values α=0.05, 0.01, to 0.001. To maximize the applicability of this study, the POD tables and graphs included in this study have been constructed with experimental scientists in mind, whose main concerns are practical considerations for designing robust and conclusive experiments (note #2). For this reason, we have determined power of detection as a function of the % change in mean lifespan, rather than as a function of the hazard ratios. Our decision was based on a biological and a practical reason. The biological reason was that many interesting experiments require the comparison of populations with different γ rather than proportional hazards (Fig. 1d and Fig. 4, below). Practically, it is conventional for most geneticists using invertebrate model organisms to study aging to report changes in mean lifespan rather than changes in hazard ratios.

In our analysis, we simulated populates with increased lifespan. As shown in Fig. 1d, the Gompertz equation allows for the generation of long-lived populations by either reducing the value of G (red graphs) or γ (green graphs) or both. Throughout this manuscript, red curves have been generated by modulating G while green curves have been generated by modulating g. Modulating G (red) causes a parallel shift of the mortality curve h(t), and thus a parallel shift in the entire lifespan curve S(t). In contrast, modulating γ (green) reduces the slope of the mortality curve h(t), and flattens the lifespan curve S(t) (Fig. 1d). We decided to first generate long lived populations by modulating G rather than γ, as this does not alter the properties of the f(t) distribution. However, as conditions that modulate γ are particularly interesting from a biological perspective as this might slow the rate of aging, we separately consider this situation later in the manuscript.

### Effect of frequency of scoring on POD

Ideal experimental planning minimizes workload without compromising the ability to collect data necessary to test the hypothesis in question. The frequency of scoring for dead animals during lifespan experiments is a major determinant of the workload of lifespan experiments. We therefore used our parametric model to determine how the frequency of scoring the number of dead animals in lifespan experiments affects mean lifespan and POD. Fig. 2a shows how the lifespan graphs changes by increasing the frequency of scoring sessions from once every twenty days to once every 12h. The further spaced scoring sessions are, the larger the error in mean and median lifespan, which is asymmetric and always increases lifespan. The asymmetry is due to the nature of the experiment: if the animal dies shortly after it was determined to be alive, its death will only become apparent during the next session. If that session is 12h or 10 days away will clearly make a difference as the animal is recorded to have died on the day death was discovered. Thus, the error will increase the apparent lifespan of the animal proportionally to the space between scoring sessions. Fig. 2b shows the percent increase in lifespan as a function of the frequency of scoring. For organisms with a relatively short lifespan, like *C. elegans*, the scoring frequency causes considerable differences in the measured mean lifespan. This effect may contribute to the large variation of N2 lifespan observed across the literature (Gems & Riddle 2000).

**Fig. 2.**
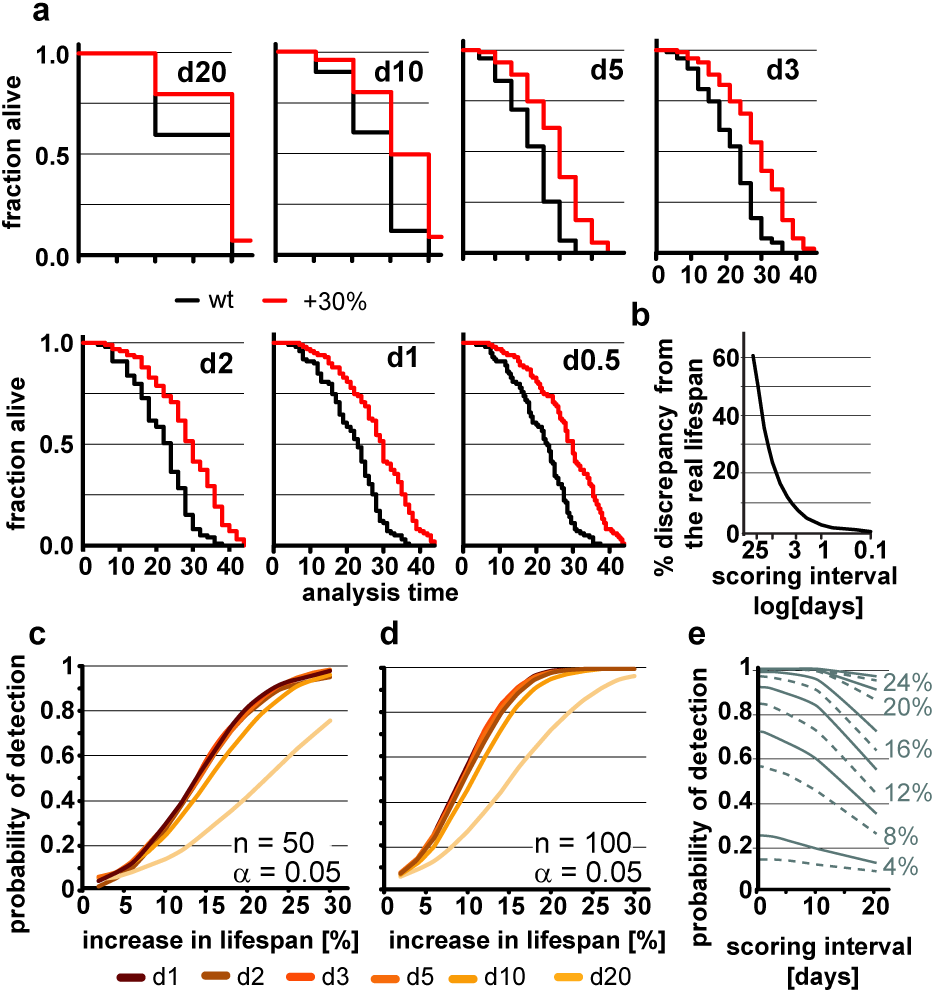
Frequent scoring of lifespan strongly affects accuracy of lifespan measurement but only modestly affects power of detection. a) Lifespan curves simulated from populations of 100 animals/cohort with a difference in lifespan of 30%. Scoring was every 20, 10, 5, 3, 2, 1, 0.5 days as indicated. b) Relationship between scoring frequency and the difference between measured and real differences in lifespan. c) POD (power of detection) plot, showing the probability to detect a significant increase in lifespan as a function of % increase using 50 animals and a significance level α =0.05. The frequency of scoring the fraction of animals alive or dead every 20, 10, 5, 3, 2, 1 of 0.5 days. Each data point was simulated 10,000 times. d) same as c but using 100 animals/cohort. e) Graphs the probability to detection as a function of scoring frequency (interval between session) for lifespan extensions ranging from 4 to 28% (n=100, α=0.05). Note that the loss in detection power is most pronounced for lifespan extensions of 8 to 16%. The labels on the right indicate the % change in lifespan and label the dashed lines. Dashed and solid lines were alternated to enhance clarity. (See supplementary table 1 for precise values)

We next used our model to determine how the scoring frequency affects POD. As mentioned above, we constructed POD plots by randomly generating 10,000 pairs of lifespan datasets for each data point, increasing the lifespan of one of the two sets by increments of 2%, and analyzed each pair by the log-rank test. To account for the difference in scoring frequencies each day of death was rounded up to the next scoring interval. For each scoring interval (e.g. every day, every 2^nd^ day) we plotted the probability to observe a significant lifespan extension as a function of the % increase in lifespan for populations with either 50 (Fig. 2c) or 100 (Fig. 2d) animals using a common significance level α = 0.05. The associated POD plots reveal two insights. First, the POD depends on the frequency on scoring dead animals (Fig. 2c-e). There was a clear difference in the ability to detect changes in lifespan when animal survival was scored every 0.5 days as compared to every 20 days. Second, the effect of increasing the frequency of scoring on POD dropped off rather dramatically. We observed relatively little change in the POD between scoring the number of dead animals every 12h or every 5 days, even though there is a 10-fold difference in workload between these two experimental approaches (Fig. 2c, d). Thus, increasing the frequency of scoring increases the accuracy of the determination of mean and median lifespan (Fig. 2b), but has surprisingly little effect on POD for changes in lifespan. Note however, that this statement holds only if test and control cohort are scored on the exact same day.

The small effect of repeated scoring on POD surprised us. We therefore analyzed the effect in more detail and asked whether increasing the frequency of scoring could improve the POD in experiments in which the number of animals assayed is low relative to the expected effect. As we will show below, a population of 100 animals detects lifespan increases from 8 to 16% in less than half of the experiments (underpowered) and thus an n of 100 is not large enough to reliably detect such effects. In underpowered experiments, frequent scoring of animals improves POD and has a bigger effect than on sufficiently powered experiments. Figure 2e shows that for lifespan extensions of 20% or more the scoring frequency has little effect on POD as the cohort of 100 animals is sufficient for robust detection. However, for lifespan extensions between 8 to 16% cohorts of 100 animals are not sufficient to reliably detect the effect on lifespan and frequent scoring becomes important as to not diminish POD further (Fig. 2e). Taken together these simulations show that the frequency of scoring of dead animals proportionally improves the accuracy by which mean and median lifespan can be determined (Fig. 2b) and that scoring frequencies are an important parameter to be considered when mean lifespans are compared between labs. They further establish that the frequency of scoring has relatively little effect on POD for appropriate sized samples but becomes important if sample sizes are low (Fig. 2c-e).

### Effect of population size on POD

The number of animals to be scored for lifespan also contributes greatly to the work required for an experimental determination of lifespan. Thus, it is preferable to test the smallest possible cohort sizes. However, it is intuitive that increasing the number of animals has a dramatic effect on how consistently changes in lifespan can be detected, which becomes already apparent in POD graphs in Fig. 2c, d. Consistent with this, we found that a 30% increase in lifespan could not reliable be detected from populations of 20 animals each in simulated experiments (Fig. 3a). We used a three-day scoring interval for these simulations (d3). Although there is a 30% difference between the two populations, this difference is detectable in only one of the two *in silico* experiments (Fig. 3b, c). Note that failure to detect the real 30% increase in lifespan in Fig. 3c is related to the control cohort (black) that lives longer than expected simply due to random fluctuations. Also note that Fig. 3b arrives at the correct result only by accident. Both, the control lifespan (black 21.2 days) as well as the increased lifespan (27.9 days) are shorter than expected. As we have shown above, counting every three days proportionally increases the lifespan due to the asymmetric error. Thus, the lifespans should be roughly 1.5 days longer than the real lifespan shown in Fig. 3a. However, due to the low number of animals, both cohort happen to be shorter lived and thus, entirely accidentally, the experiment arrives at the true increase in lifespan of 30%. The advantage of doing these experiments *in silico* is that we know for certain that these results arise through random fluctuation and not due to environmental differences or inadequate technique by one of the researchers. These graphs illustrate that random fluctuations preclude the ability to reliably detect a 30% change in lifespan when the cohort size is 20 animals or less.

**Fig. 3.**
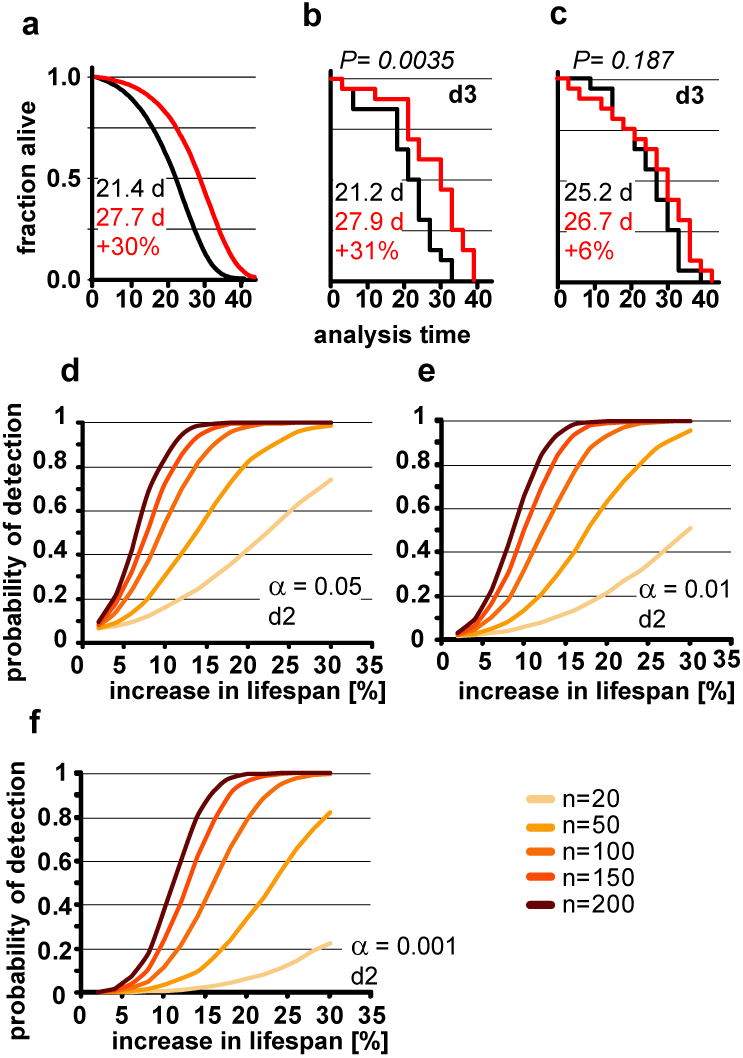
Population size has important effect on power of detection. a) Parametric model as in Fig 1 (left) and Kaplan-Meier plots of two independently simulated lifespan experiments (b, c) using n=20 for wild-type animals (black) and a perturbation that increases lifespan by 30% due to a change in G (red). Scoring frequency indicated in the upper right corner, P value indicated above each graph. Mean lifespan in days and % change for both cohorts is indicated on each graph. d) Power of detection plotted as a function of % increase in lifespan for α=0.05 and a 2-day scoring frequency (d2) in determining the fraction of animals dead/alive. Cohort sizes ranging from 20 to 200 animals. Lifespan was increased by modulating the parameter G in the Gompertz equation. e) same as d) but α=0.01. f) same as d) but α=0.001. For b-d each data point was simulated 10,000 times. (See supplementary table 2 for precise values)

Although lifespan experiments with only 20 animals in each cohort may be somewhat of an extreme example, our simulations illustrate the difficulty in interpreting underpowered experiments. In our example, we know that there is a true increase of 30% in the mean lifespan as detected in Fig. 3b. However, without this *a priori* knowledge we could not distinguish whether the difference between Fig. 3b and c arose through environmental differences, inadequate experimental skill, or random fluctuation. In short, if an experiment is underpowered, random fluctuations cannot be distinguished from real biological effects.

In order to determine the number of animals required to detect changes in lifespan of various magnitude, we used our model to calculate how the POD changes a function of % increase in lifespan for different population sizes (Fig. 3d-f). For each of these experiments, we maintained the significance levels a as well as the scoring frequency constant (every 2 days). As expected, these POD graphs show that for any given change in lifespan, the POD is higher with larger cohorts of animals. The POD graphs further reveal that the number of animals required to reliably detect smaller changes in lifespan require substantially more animals than are generally used for lifespan experiments. For example, to detect a true 12% increase in lifespan at a significance level α=0.05 in at least 4 out of 5 experiments (>80%) requires 150 animals for each cohort (control and test cohort) or a total of 300 animals. If we use a more stringent significance level of α=0.01 will require 200 animals in each cohort, a total of 400 animals. Note that the number of animals required will increase further if the animals have differences in genetic background or if there are additional factors increasing the variance which are not part of our model (we have included a list of common confounding factors in the supplementary material). Based on these data, a sizable number of publications reporting increases in lifespan of 10-15% are seriously underpowered (citations intentionally omitted, see note #1 in Supplementary materials). This does not automatically mean that these results are all erroneous, but that a definitive conclusion cannot drawn from the lifespan experiments alone, due to the small cohort sizes.

For the experimental scientist, the POD graphs shown in Fig. 3 provide a reasonable guide of how many animals are required to reliably detect a given change of lifespan for *C. elegans* experiments. However, we want to stress that these graphs constitute a best-case scenario. Any additional variation resulting from experimental conditions is more likely to increase the number of animals required (see supplementary data). The only scenario that would reduce the numbers shown in Fig. 3 and in the supplementary POD Table, would be a tighter distribution in the lifespan data. It is possible that lifespan data from some labs show a slightly tighter distribution than the one in our model for two reasons. First, to some degree, our model incorporates researcher to researcher variation as the original dataset was derived from lifespan data determined by two researchers, Second, censoring early deaths will tighten the death time distribution f(t). The data used for our model have a censoring rate of less than 1%, and were censored only at the end, but not at the beginning of the experiment. However, even if the distribution achieved by others is slightly tighter, which should be verified by deriving γ from a data set of several hundred control animals, the reductions in sample size are likely going to be minor. Therefore, we suggest that the numbers presented should be read with an “at least” in mind. Together, the data in Figs 2 and 3 indicate that the optimal experimental design to verify the reproducibility of a lifespan extending effect should include more animals in each cohort with a longer interval between scoring session to most efficiently maximize the POD.

### Effect of altered hazard distributions on POD

Our parametric model is based on the Gompertz equation, so lifespan can not only be modulated by modulating G, as we have used in the previous experiments, but also by modulating γ (Fig. 1). Extending lifespan by changing γ changes the pace by which mortality increases (Fig. 4a). Interventions (e.g. mutations, compounds) that slow the pace of age associated mortality, compared to wild type animals, and thus show a different γ value, are of great biological interest. We therefore explored how the POD is affected in experiments involving cohorts with different γ values.

**Fig. 4.**
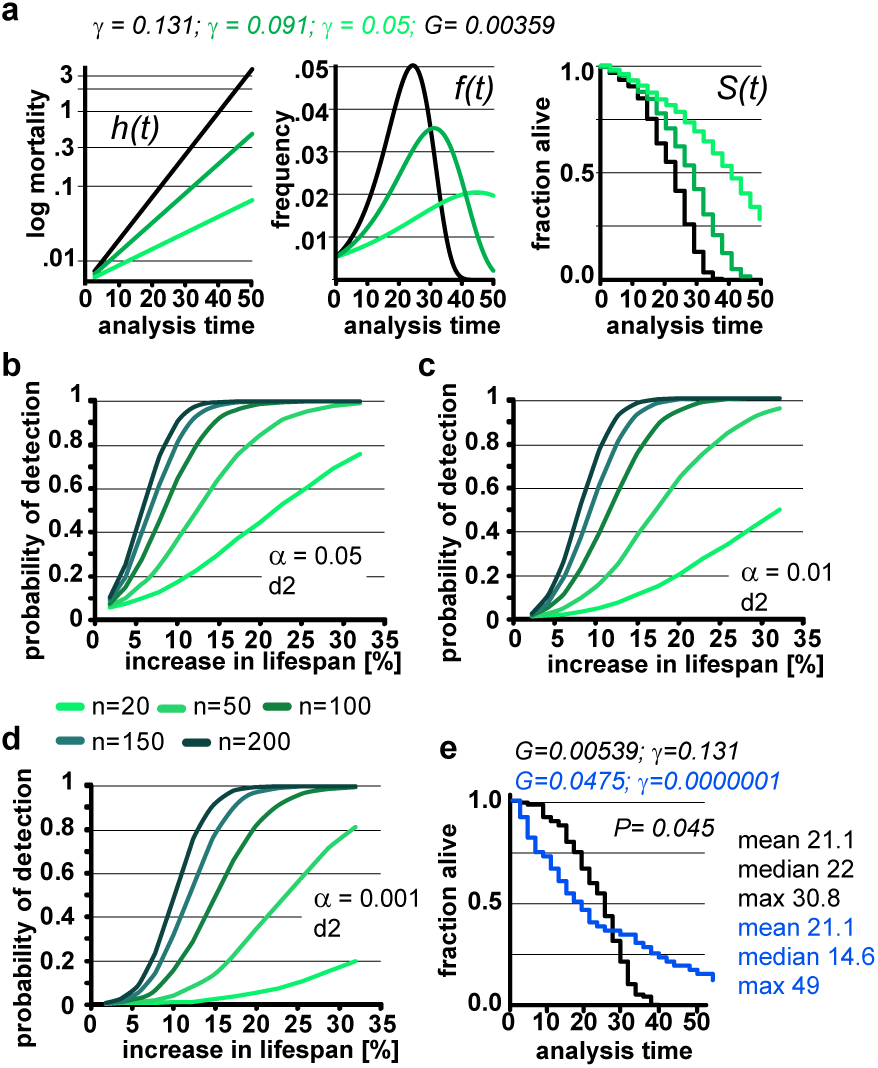
Lifespan extensions caused by changing γ values has little effect on POD. a) Demonstration of how changes in γ (green) affect the mortality curve h(t), frequency distribution f(t), and survival plots S(t). n=100 animals, scoring frequency every 2 days. b) Power of detection as a function of magnitude of % change in lifespan caused by changes in γ. Each curve is for a different cohort size, from 20 to 200 animals. α=0.05. c) same as b, but α=0.01. d) same as b, but α=0.001. For all curves, scoring was every two days (d2). Each data point was simulated 10,000 times for b-d. e) Example on how simultaneous changes in G and γ introduce difficulty in interpreting lifespan data. Lifespan curves were simulated for populations of 100 animals scored every 2 days. The two populations have the exact same mean lifespan (21.1 days), but different median lifespans (22 days, 14.6 days) and maximum lifespans (30. 8 days, 49 days). The control (black) population lives longer with respect to the median lifespan, whereas the blue population lives longer with respect to the maximum lifespan. The values for G and g are indicated for both populations above the graph. (See supplementary table 3 for precise values)

Changing γ alters the density function, f(t) and lifespan curves, S(t) (Fig. 1). In practical terms the density function f(t) is the normalized version of a histogram plotting the number of animals that died at a given day (see supplementary data). Higher γ values will cause a narrow f(t) distribution, as all the animals die within a short window of time. As can be seen in Fig. 4a, decreasing γ (green) widens the f(t) distribution and cause the animal to die of over an extended period. Decreasing γ also reduces the slope of the lifespan curve S(t), which is commonly considered to represent a decrease in the rate of aging (Gems *et al.* 2002). However, another interpretation is that γ is a measure of stochastic variation in lifespan, as reflected in the f(t) distribution. In this interpretation, the apparent rate of aging in a population is, somewhat counterintuitively, related to stochastic variation in lifespan. Although these are interesting considerations, as we noted above, the Gompertz model is not a mathematical model based on a hypothesis how aging works, but a convenient equation that describes survival time data.

We used our parametric model to simulate lifespan curves by progressively lowering the values of γ to extend lifespan (Fig. 4b-d). The POD curves generated in this experiment are surprisingly similar to POD curves we calculated in experiments where lifespan changes result from changing G (compare Fig. 4b-e with Fig.3 b-d and supplementary POD table). As an experimentalist, in which it is impossible to predict whether a given perturbation will influence G or γ, this is quite reassuring. We conclude that, for practical purposes, POD for experimental design can be determined without consideration of whether lifespan is extended by modulating G or γ.

Because the log-rank test was unexpectedly resilient to changes in γ, we explored how well the log-rank test would perform under extreme circumstances. In this test, we changed both γ and G to construct an example in which two cohorts have the exact same mean lifespan of 21.1 days but entirely different density distributions f(t) and, consequently, very different lifespan curves S(t) (Fig. 4e, blue). Although both populations have the same mean lifespan (21.1), the median lifespan of the control (black) cohort is longer (22 days vs 14.6 days), whereas the maximum lifespan of the green cohort is longer (30.8 vs 49 days). By log-rank analysis, the difference between these lifespan curves is significant (p= 0.045, n= 100) favoring the cohort with the longer maximum lifespan. This seems biologically sensible as most people would weigh the greater maximum lifespan as more important than the shorter median lifespan. Our POD tables do not account for such extreme cases as the tables are based on changes in mean lifespan. Judging from the current literature, such cases seem rare if they exist at all, but it is important to know that at least theoretically cases exist in which current methods will break down.

### Examples of how power of detection affects reproducibility

The motivation behind this study was to determine the reproducibility of results under ideal conditions, using the current practices. In the examples below, we demonstrate how inadequate POD considerations can have practical implications on the interpretation of lifespan data and their reproducibility.

#### Example 1: Effect of POD on lab to lab reproducibility

The framework we have established thus far allows us to evaluate reproducibility with random variation being the only sources of variation. The POD graphs shown in figures 2-4 plot the probability to get a significant result in a single experimental trial. However, most lifespan assays are, appropriately, repeated multiple times to test for the effect of compounds, mutations, or RNAi on lifespan. Nevertheless, when replicate experiments are conducted with underpowered cohorts, reproducibility will seriously be reduced.

Consider the situation where a group or researchers identifies a compound that has a modest effect to increase lifespan (e.g. 12%). To validate this observation, the researchers retest the compound in triplicate with cohorts of 50 animals in both the control and treated group, scoring the fraction of animals alive and dead animals every 2 days. They also send the compound to a second lab, which runs the same experiment in triplicate. The POD table (Supplementary Table) indicate that with n=50 and α = 0.05, a 12% increase in lifespan will be detected in 41% of all trials. Thus, there is a 0.5% chance (0.4^6^) that both labs will confirm the (real) effect of the drug on lifespan in all three experiments. In this situation, the underpowered experiments are likely to result in a false negative conclusion.

This example raises the question of what should be considered a “reproducible” result. A more relaxed and probably reasonable definition of reproducibility would be not to require that the effect on lifespan to be observed in all 6 trials, but only in at least 2 successful trials in each lab. This relaxed definition of reproducibility would raise the probability of reproducing the result to ~15%. This is still far from a robust result: In over ~45% of the cases the two labs will obtain contradictory results or worse, wrongly agree that that the compound does not extend lifespan (~40%). Increasing the number of animals has large effects on POD (Fig. 3), but even if both labs had used 100 animals in drug and control cohort the chance that both labs would have obtained a significant extension in all six trials would still be a mere 11.5% (POD = 69.7% per experiment). Using the relaxed definition of reproducibility (at least two or more successful trials in each lab) would raise the chance of reproducibility to 61%. Based on our experiments, to have a ~90% chance that both labs would detect the real change in lifespan in 2 or more of the three experiments would require at least 150 animals in each cohort, a minimum of 1800 animals (n=150, 2 x (control & compound) x3 trials x 2 labs= 1800). These cohort sizes are far bigger than those generally published. Note that in this example the failure to reproduce the result has nothing to do experimental skill or sloppiness but is purely the result of the variation inherent in lifespan data. On the contrary, these are best case scenarios, with perfect conditions, and identical perfect scientists performing the experiments, and still the chances of reproduction are low.

#### Example 2: Screening for compounds that can increase lifespan

A real example of how underpowered experiments can lead to unexpected results can be observed in one of our own studies. Screening for compounds that extend lifespan we consistently and non-intuitively identified more compounds that extended lifespan by 10-20% (24) than compounds that extended lifespan by 1-10% (4) (Fig. 2a in (Ye *et al.* 2014)). Our experimental design involved 80 to 100 animals, and to be considered a hit, a compound had to reproducibly extend lifespan across multiple experiments. However, 80 to 100 animals are insufficient to repeatedly detect lifespan extensions from 1-10%. Thus, many compounds with real but small effects were removed from our list of hits because the POD of our experimental design made it unlikely to repeatedly observe a significant result. Considering the POD of these experiments, we can conclude that the low number of compounds that extended lifespan by 1-10% was not the result of some mysterious biology but the direct result of the experimental design.

Examples 1 and 2 both illustrate the difficulty in reproducing small lifespan extensions. However, more encouraging is that the odds start to improve dramatically at increases of +20% or more. In example 1, even using only 50 animals in each cohort, the chances for reproducibility between two labs are 84%, and for effects sizes of +30% it is close to 100% under ideal conditions. This result highlights that POD becomes increasingly important for lifespan extensions below 20%. This example also highlights another caveat of how lifespan results are communicated: reporting lifespan extension by citing the best % increase in lifespan, generally published as an “up to X% increase in lifespan”, is counterproductive to reproducibility. Creating the impression that the highest increase in lifespan is equivalent to the mean increase in lifespan by using fuzzy and unclear language will lead to underpowered experiments by others trying to reproduce these results. This will result in a reproducibility problem, not because the intervention doesn’t work, but because it doesn’t work as well as advertised.

#### Example 3: POD considerations for epistasis analysis

This last, and in our view, most serious problem is caused by underpowered studies in epistasis analysis. Let’s again assume researchers have identified a mutant that extends lifespan by 12% after having diligently outcrossed the strain. They compared the lifespan of this mutant to N2 in six independent experiments using 50 animals in each cohort. They only observed a significant effect in three out of these 6 experiments (chance to detect 3 or more =47%). Pooling the data, however, the effect on lifespan became highly significant, convincing them (correctly, in this situation) that the mutant is long-lived.

To determine whether the longevity effect acts through the insulin-signaling pathway, the experimenters next decided to conduct experiments to determine if the increased lifespan depends on the FOXO transcription factor DAF-16. To avoid genetic background effects, the researchers used *daf-16* RNAi in 4 independent trials with 50 animals in each cohort. In truth, and unknown to them, the longevity effect is completely independent of *daf-16*. Unfortunately, in their experimental setup the probability to miss the 12% increase in lifespan in <u>*at least*</u> 3 of the 4 trials is 46%. This does not even take into consideration variation resulting from RNAi itself. Thus, the chances arriving at the correct result by conducting a series of 4 trials using their experimental design are only marginally better than that of a coin toss, despite the substantially greater effort involved. In nearly half of the experimental series, the longevity effect will appear to be *daf-16* dependent because no lifespan extension will be detected in at least 3 out of 4 experiments. If a lifespan extension is observed in only one of the 4 experiments, it is easily explained away by the occasional inefficiency of RNAi. The only indication that the lifespan extension might in truth be *daf-16* independent will be uncovered by pooling the data (n=200). Pooling the data in which case a lifespan extension will become apparent in 94% of the experimental series. However, this indication will only become apparent if the variability between the experiments is minimal. *In silico* pooling is an effective strategy as there is no inter-experimental variation; however, pooling real-life data is highly problematic due to the variability between experiments, which will increase the overall variation obscuring any real effect further, or causing non-existing effects. Even if pooling is successful, no sensible reviewer will accept a claim of *daf-16* independence if 3 out of 4 experiment show no significant lifespan extension.

### Conclusions

In this study, we generated a parametric model of *C. elegans* lifespan from a large cohort of wild-type lifespan data. We used this model to demonstrate the importance of considering POD in experimental design to measure the effect of genetic or pharmacological perturbations of lifespan. Based on our data, we provide tables to facilitate the planning of rigorous lifespan experiments. Avoiding insufficiently powered studies will improve the reproducibility of lifespan experiments within and between labs, thereby enhancing the efficiency of research in this important field.

## Experimental Procedures

*The experimental method used to generate the lifespan data of 5,026 N2 animals which formed the basis of the parametric model can be found in Solis and Petrascheck (Solis & Petrascheck 2011).*

### Generation of a parametric survival-time model

The model used in this publication is similar as we have used previously to model some of our lifespan screens (Ye *et al.* 2014). It is parametric survival-time model based on the Gompertz equation. Using this survival-time model we generated artificial datasets using Monte-Carlo simulations. Each data point was simulated 10,000 times. Simulating a parametric model allowed us great flexibility in changing experimental practices. It may be possible to derive the exact equations without the need of simulations but our approach avoided some rather intimidating equations, some of which we were admittedly unable to solve.

#### Definitions

T: non-negative random variable denoting the time to a death event

F(t): cumulative distribution function: *F*(*t*) = Pr (*T* ≤ *t*)

S(t): survivor function, lifespan curve: *S*(*t*) = Pr (*T* > *t*) = 1 − *F*(*t*) *and f or S*(0) = 1

Gompertz equation: *h*(*t*) = *G* _*_ *e*^(*t*\**γ*)^

Equations to derive the quantile function *Q*(*u*) = *F*(*u*)^−1^ necessary to simulate lifespan data with the same (or altered) characteristics as the lifespan data obtained from the 5,026 N2 control animals.

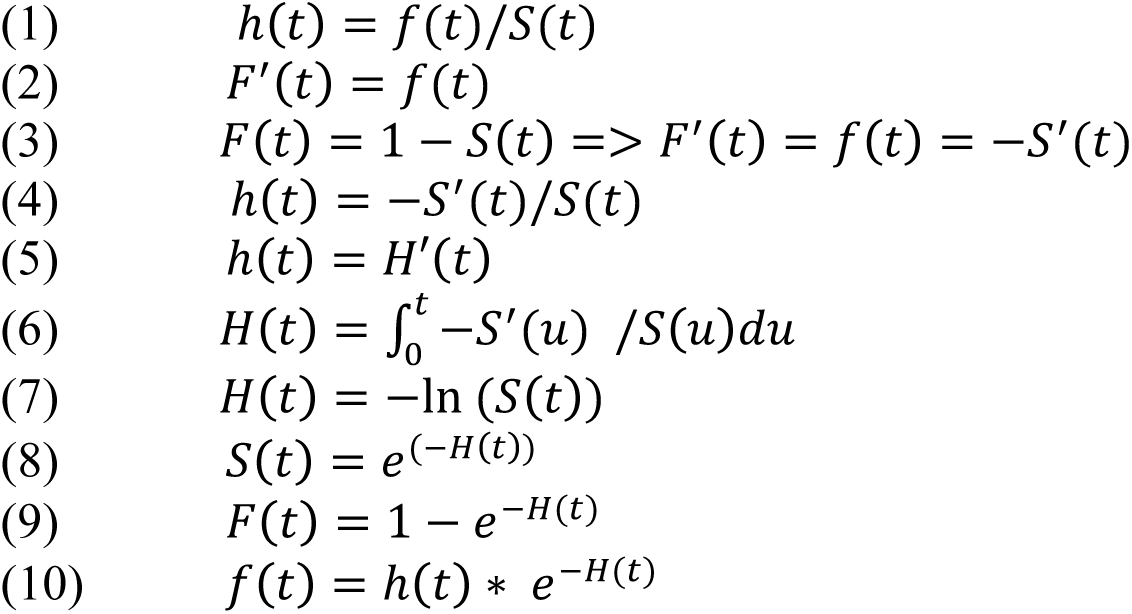

after determining the relations of S(t), F(t) and f(t) to h(t) we can introduce the Gompertz equation h(t).

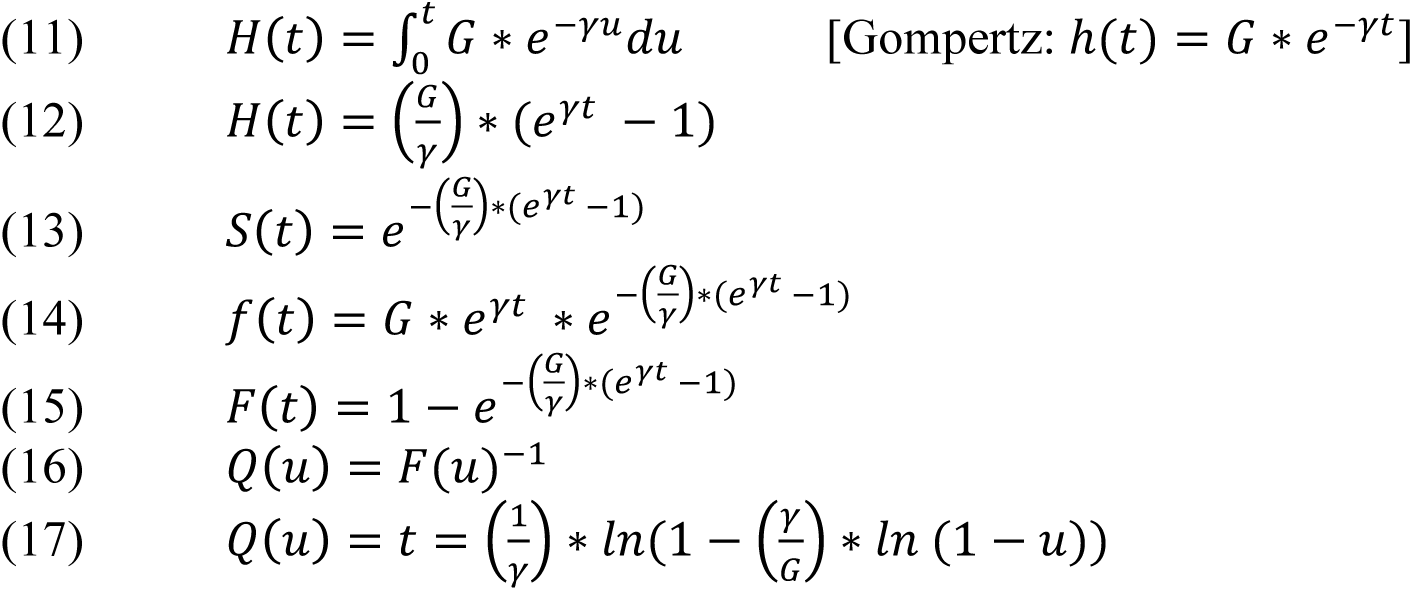

Maximum likelihood estimation was performed using the STATA streg command to estimate the parameter γ and subsequently G from the survival-time data of the 5,026 control animals. To simulate the survival-times for control and long lived animals, the quantile function, defined as the inverse of the cumulative distribution F(t) for the Gompertz was obtained (equation 17). Monte Carlo simulations were then used to artificially generate datasets using randomly generated numbers for *u* (0 ≤ *u* < 1) resulting death times *t* that show the same distributions as real *C. elegans* data using Q(u) (equation 17). Q(u) in combination with a random number generator generating values for u (0 ≤ *u* < 1) can be used to simulate lifespan data for any species by determining γ and G using regression analysis from real data.

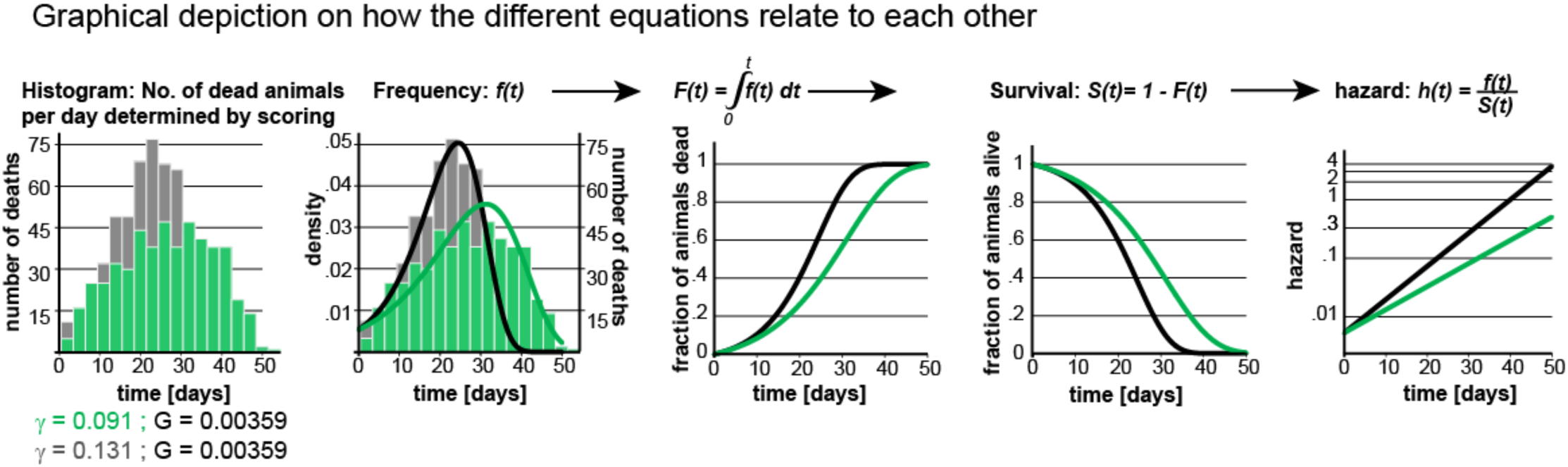

#### Construction of POD plots

For each data point in each curve, we generated 10,000 paired datasets consisting of a control and test population and evaluated potential statistical differences using the log-rank test. The power of detection (POD), or probability of detection, was determined as the fraction of times a true lifespan extension was detected for a given significance level a. For the POD curves shown in Fig. 2 we maintained a constant number of animals n=50 or 100 and a significance level α = 0.05 but altered the frequency of scoring by rounding up death times to the next scoring session. To plot each curve, we increased the lifespan in increments of 2% (by changing G), each time simulating 10,000 datasets to determine the probability of detection by the log-rank test (Mantel, Haenszel, 1959 version). We then plotted the probability of detection for different scoring intervals as a function of % increase in lifespan.

For the POD curves shown in Fig. 3 we maintained a constant scoring interval of 2 days, a significance level a of 0.05 (b), 0.01(c), 0.001(d) and plotted the probability of detection as a function of % lifespan increase for different number of animals. As before, we simulated 10,000 datasets to determine the probability of detection for each lifespan increment of 2% (by changing G).

The POD curves shown in Fig. 4 were constructed as the POD curves in Fig. 3 except this time we modulated γ instead of G to extend lifespan.

Probabilities in examples 1 and 3 were calculated by generating the entire probability tree diagrams and by subsequently adding up the probabilities of successful and unsuccessful outcomes, as defined in the example.

## Acknowledgements

We are grateful to Greogry Solis and Maria Carreterro for critical comments. This work was supported by NIH grants R01 AG044378 (DLM), R01 ES024958 (DLM) and DP2 (MP) and the Glenn Foundation (DLM & MP). DLM and MP were New Scholars in Aging of the Ellison Medical Foundation.

## Supporting Information listing

POD tables

Notes

Sources of Variation and Limitation

## Supplementary material

### POD Tables: See excel, POD tables

Note #1: There have been many public and private disputes regarding lifespan experiments. The origins of these disputes are often unclear. In many of these disputes POD issues may have contributed or even be the underlying reason or may have not played a role at all. We therefore have not included any citations regarding these controversies. We considered citing controversies counter productive in the absence of a thorough analysis of each individual case, which we were not prepared to conduct.

Note #2: The intended reader of this manuscript are experimental scientists that are users of statistics and not statisticians. We have written this paper avoiding mathematical formalism as much as possible focusing extensively on the practical consequences and implications. As a result, we occasionally over-simplify certain aspects or deviated from standard practices. For example, our POD graphs are plotted as a function of effect size, a practice that can be criticized on various grounds, but which is the most user-friendly way of presenting POD curves.

### Sources of variation and limitations

As our parametric model is derived from actual data some forms of variation are accounted for while others are not. In this section, we will shortly elaborate which sources of variation are taken into account and which ones are not, and thus will further increase the number of animals necessary for a sufficient POD.

*Different researchers:* In our experience, when different people repeat the same experiment the same result is generally detected. However, if the number of animals is large enough (n=200-500), there may be statistically detectable differences in the measured lifespan of control animals between researches (up to 5%). Direct comparison of the scoring data of 3 different researchers scoring the same cohort suggests that some experimenters tend to consistently measure longer lifespans than others. The lifespan data of the 5,026 animals used for deriving our parametric model contains data generated by two different researchers, and thus researcher to researcher variation is built into the model. We also propose that very small changes in lifespan (1-10%) should be viewed with extra skepticism in light of this source of variation.

*Plate to plate variation*: Especially in experiments with large cohort sizes, the animals are distributed across different plates. There can be minor differences in lifespan between plates. This source of variation is included in our parametric model, as the original lifespan data of 5,026 animals used to derive the model were distributed across 64 different plates.

*Undefined media and bacteria:* The lifespan data of the 5,026 animals used to derive the parametric model were all fed bacteria from a single culture, and lifespan experiments were conducted in liquid medium with the only undefined substance being bacteria but no agar or bacto peptone. Agar is a plant extract and bacto peptone is made from beef and pork trash like brain, ovaries, gut and testes (Kamekura *et al.* 1988). Dependent on the origin these undefined substances contain hormone residues such as bile acids, estradiols, and other signaling hormones that could influence lifespan. While these ingredients are unlikely to affect single experiments, if control and test animals are cultured on the same batch, they are likely to increase variability between labs that use different lots. Multi-lab experiments are probably best conducted with ingredients with identical lot numbers to minimize variation and thus enhance reproducibility.

*Automation:* Recent advances in hardware have resulted in the generation of more high-throughput lifespan data using image based automation methods (Xian *et al.* 2013; Stroustrup *et al.* 2016; Zhang *et al.* 2016). These automated methods increase throughput for lifespan experiments and will increase cohort sizes thus generally increasing reproducibility. However, these methods generally require a “data cleaning step” to remove spurious data which can introduce systematic errors affecting POD limits. We recommend that every automated method should be compared carefully compared to human counting, and that a coefficient of variation or similar measure be included when these data are published to enable appropriate comparisons to human data.

